# SCARECROW maintains the stem cell niche in Arabidopsis root by ensuring telomere integrity

**DOI:** 10.1101/2021.10.25.465752

**Authors:** Bingxin Wang, Xiaowen Shi, Jingbo Gao, Rui Liao, Jing Fu, Juan Bai, Hongchang Cui

**Affiliations:** State Key Laboratory of Crop Stress Biology for Arid Areas and College of Life Sciences, Northwest A&F University, Yangling, Shaanxi 712100, China; Department of Biological Science, Florida State University, Tallahassee, FL 32306, USA

## Abstract

Stem cells are the ultimate source of the cells in various tissues and organs, and thus are essential to postembryonic plant growth and development. SCARECROW (SCR) is a plant-specific transcription regulator well known for its role in stem-cell renewal in plant roots, but the mechanism by which SCR exerts this function is still unclear. To address this question, we carried out a genetic screen for mutants that no longer express SCR in the stem-cell niche in the Arabidopsis root, and one of the mutants is characterized herein. Using marker-assisted mapping, whole genome sequencing, and complementation tests, we pinpointed the causal mutation in this mutant at *TEN1*, which encodes telomere-end protecting factor. By sequence alignment of TEN1 homologs in a wide range of eukaryotes, we identified two novel motifs. The importance of these motifs was examined through site-directed mutagenesis of a conserved amino acid, and through complementation tests, which showed that G100 is required for TEN1 stability. Interestingly, we found that *TEN1* expression was dramatically reduced in the *scr* mutant. Two components in the same protein complex as TEN1, STN1 and CTC1, were found to be also dramatically downregulated in *scr*, as well as telomerase. Further studies showed that loss of STN1, CTC1 and telomerase also caused defects in the root stem cells. In line with these findings, the *scr* mutant was hypersensitive to DNA damage reagents such as Zeocin. These results together suggest that SCR maintains root stem cells by promoting expression of genes that ensures genome integrity.

**One sentence summary:** This study reveals a role for the root development regulator SCARECROW in maintaining the expression of telomere-end protecting factors and a connection between genome integrity and stem-cell maintenance in Arabidopsis root.

## Introduction

Postembryonic growth and development in higher plants relies on two populations of stem cells, one at the root tip and the other at the shoot tip. Located at the center of the root tip, the quiescent center (QC) is the ultimate source of all root cells and is therefore considered the very stem cell of the root (Fig 1A). The QC is surrounded by cells that also have stem cell properties but give rise to various cell types, and together these cells constitute the so called stem cell niche (SCN) (Benfey and Scheres, 2000), in which the QC acts as the organizing center (Dolan et al., 1993; van den Berg et al., 1997). Daughter cells of the SCN are mitotically much more active, forming a zone of dividing cells named the root apical meristem (RAM), which is adjoined by the elongation zone (Fig 1A). Due to its critical importance in root growth, the SCN has been be the subject of extensive research, and many factors involved in SCN maintenance have been identified in the model plant *Arabidopsis thaliana*, such as the *PLETHORA* family genes (Aida et al., 2004); *WOX5*, a *WUSCHEL* family transcription factor specifically expressed in the QC (Sarkar et al., 2007); and TCP family members TCP20 and TCP21 (Shimotohno et al., 2018).

**Fig 1.**
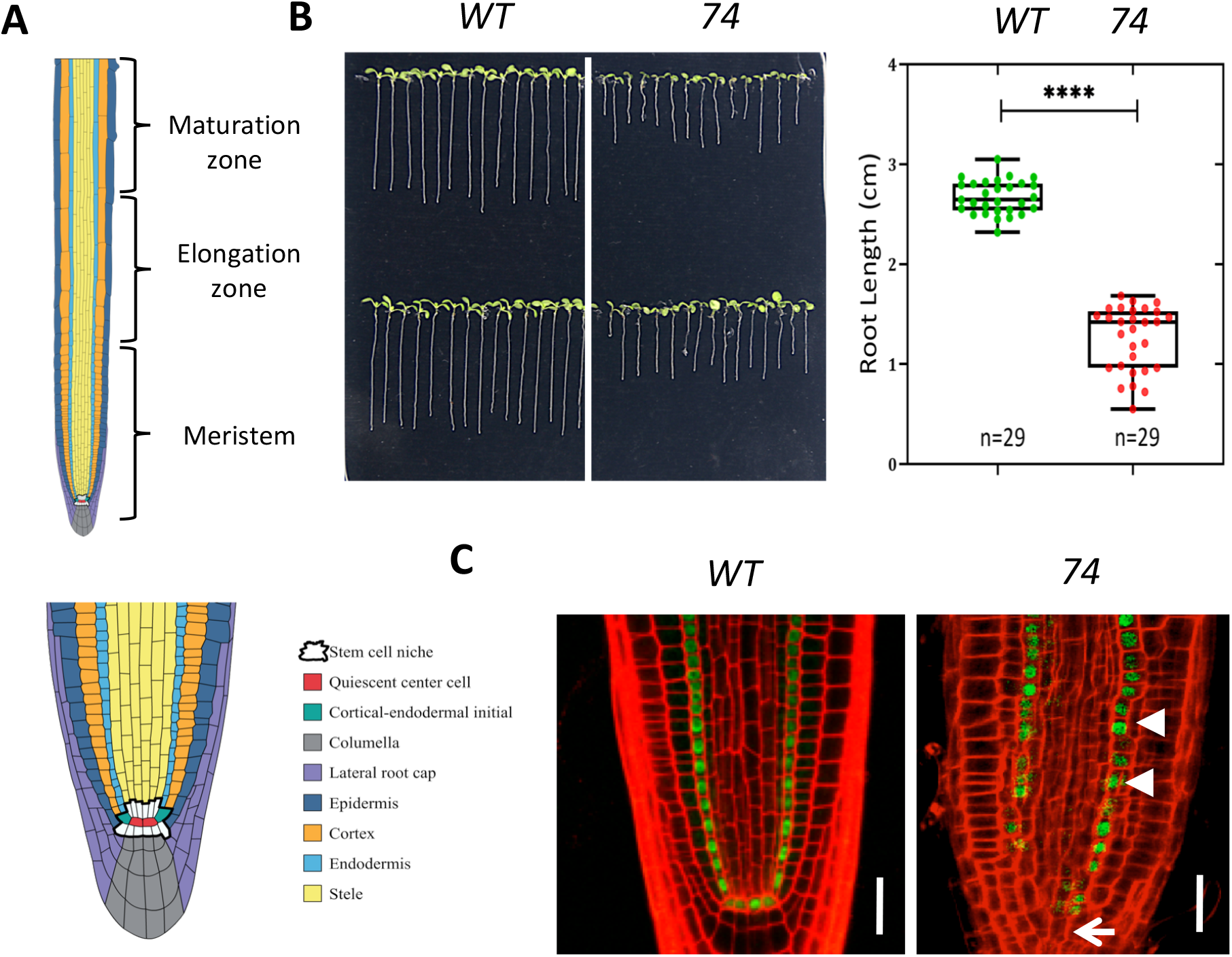
Mutant 74 is defective in root stem cell maintenance. A. Diagram of the different developmental zones (upper panel) of Arabidopsis root and cell types (lower panel) in the apical root meristem. B. Root length of 8-day-old seedlings. Representative image (B) and box plot (C). **** represents *p* < 0.0001, *t*-test. C. Confocal microscopy image showing the expression pattern of the *SCRpro::GFP-SCR* transgene in the roots of wild type and mutant 74. The seedlings were seven days old. Bar = 20 μm.

Telomere protective factors have also been shown to play a pivotal role in stem-cell maintenance. In a genetic screen for mutants with compromised root growth, Hashimura et al. identified a mutant that has a defective SCN and a short root phenotype, which they named *meristem disorganized* (*mdo1*) (Hashimura and Ueguchi, 2011). MDO1 turns out to be TEN1, which is a component of the CST complex that binds to and protects the telomere (Leehy et al., 2013). The *ten1/ mdo1* mutant is hypersensitive to DNA damage reagents, especially cells in the SCN (Hashimura and Ueguchi, 2011). Severe growth retardation and telomere defects were also observed in mutants for CTC1 and STN1, which are components of the CST complex (Song et al., 2008; Surovtseva et al., 2009), suggesting that these two protein may also have an importance role in stem-cell maintenance.

SCR is a member of the plant-specific GRAS family of transcriptional regulators with a pivotal role in stem cell maintenance and radial patterning in the root of the model plant *Arabidopsis thaliana* (Di Laurenzio et al., 1996). In the *scr* mutant, QC is lost and SCN becomes disorganized, resulting in a short root phenotype (Sabatini et al., 2003). In addition to its important role in stem-cell maintenance, SCR also plays an essential role in radial patterning. Compared to the wild type, which produces two layers of ground tissue – the endodermis and cortex – through an asymmetric cell division of the cortex/endodermis initial (CEI) cells (Fig 1A), the *scr* mutant has only a single layer of ground tissue (Di Laurenzio et al., 1996). Because this mutant cell layer has characteristics of both endodermis and cortex, it has long been thought that SCR is required only for the asymmetric division of the CEI. However, a recent study showed that SCR is required for endodermal specification as well, whereby it acts redundantly with SCL23, a close homolog to SCR (Long et al., 2015a).

The mechanism by which SCR regulates ground tissue patterning in *Arabidopsis thaliana* has been largely elucidated. Acting upstream of SCR is SHR. However, SHR is expressed in the inner tissue of the root – the stele (Helariutta et al., 2000), unlike SCR, which is expressed specifically in the QC and endodermis (Di Laurenzio et al., 1996). The SHR protein is a mobile molecule able to move into the neighboring cells –the QC and endodermis (Nakajima et al., 2001), where it forms a complex with SCR, enhances SCR expression through a feed forward loop, and specifies the endodermal cell fate (Cui et al., 2007; Sozzani et al., 2010). In the meantime, SHR gets trapped in the endodermis and QC as a result of physical interaction with and nuclear retention by SCR, thus defining a single layer of endodermis (Cui et al., 2007). This mechanism also requires some members of the IDD family of transcription factors, which physically interact with SHR and SCR as well (Long et al., 2015b; Moreno-Risueno et al., 2015; Welch et al., 2007).

Despite its pivotal role in stem cell maintenance, how SCR executes this role is still poorly understood. It is clear that expression of SCR in the QC is required for maintenance of the SCN, but whether and how SCR exerts this role has been unclear (Sabatini et al., 2003). Through transcriptome analysis, Moubayidin et al. found that cytokinin signaling is elevated in the *scr* mutant; when a cytokinin signaling was blocked by mutation in *ARR1*, the short root phenotype of the *scr* mutant is largely rescued owing to a longer RAM (Moubayidin et al., 2013), suggesting that SCR promotes root growth partially by suppressing cytokinin biosynthesis or signaling. The SCN in *scr* was also alleviated by *arr1*, as indicated by the restoration of *WOX5* expression (Moubayidin et al., 2013). However, the rescue is likely to be an indirect effect of ARR1 because ARR1 is mainly expressed in the meristematic cells outside of the SCN. Moreover, the root of the *arr1 scr* double mutant is still significantly shorter than the wild type, suggesting the existence of other mechanisms. Recently Shimotohno et al. showed that SCR is able to form a complex with TCP20 and PLT and together they activate the expression of WOX5, thus maintaining the SCN (Shimotohno et al., 2018). Although the *tcp20* mutant enhances the SCN defect in *scr*, it alone does not disturb the SCN and root growth (Shimotohno et al., 2018). In a recent study we also showed that SCR has a role in promoting cell elongation and meristematic activity by maintaining redox homeostasis, but this does not seem to be relevant to SCN (Fu et al., 2021). These studies suggest that additional mechanisms exist for SCR in SCN maintenance.

The notion that SCR functions differently in the SCN and RAM is supported by other studies. Using a promoter-bashing approach, Kobayashi et al revealed that the SCR promoter contains two cis-regulatory modules with distinct functions: one for QC expression, and the other for endodermal expression (Kobayashi et al., 2017). The factors that bind to these cis-regulatory motifs cannot be SHR, not only because SHR is present in the QC and endodermis, but also because it does not have a DNA binding domain. Members of the IDD family of transcription factors could fit this role, but most IDD genes studied so far appear to be solely involved in ground tissue patterning (Long et al., 2015b; Moreno-Risueno et al., 2015; Welch et al., 2007). The only IDD factor known to be essential to stem cell maintenance is JKD, but the *jkd* mutant has only slightly shorter root than the wild type (Welch et al., 2007), which suggests the existence of additional factors that maintain the SCN.

Understanding the mechanism by which SCR maintains SCN necessitates the identification of factors that act upstream and downstream of SCR. We therefore have designed a genetic screen that allows us to identify mutants that affect SCR expression in the QC, but not in the endodermis. Several mutants fitting this criterium were obtained, and characterization of one of them is described in this report. Through marker-assisted crude mapping, whole genome resequencing, and functional complementation tests, we located the gene with the causal mutation *TEN1*, which encodes a component of the CST complex that protects telomere ends (Shay and Wright, 2019). Interestingly, we found that TEN1 expression was dramatically reduced in the *scr* mutant. The other components of the CST complex, STN1 and CTC1, were also affected in *scr*, as was telomerase. These findings along with other relevant experiments suggest that SCR maintains the stem cell niche by ensuring telomere integrity. In addition, we have identified two evolutionarily conserved motifs in TEN1 and provided evidence that one of them has an essential role in TEN1 function.

## Results

### 1. A genetic screen for genes involved in the SCR pathway identified a mutant defective in root stem-cell maintenance

To identify factors that are involved in the SCR developmental pathway, we carried out an EMS (ethyl methanesulfonate) mutagenesis with seeds homozygous for a transgene expressing a GFP-SCR fusion protein under the control of the *SCR* promoter in the Columbia-0 (Col-0) background (*SCRpro:GFP-SCR, scr-1*). From 10,000 mutagenized seeds several mutants that no longer express GFP-SCR in the QC were obtained. One of these mutants, numbered 74, was chosen for further characterization in this study due to its apparent root growth defect (Fig 1B). In this mutant, GFP signal was lost not only in the QC but also in some endodermal cells close to the QC (Fig 1C), which suggests that the mutation affects the stem cell niche (SCN). Consistent with this observation, we found that cells in SCN became disorganized (Fig 1C). In addition to the SCN and root growth defects, the mutant also displayed abnormal phenotypes in the shoot, such as short stature, fascinated inflorescence, extra branching, and clustered flowers (Fig S1, A and B). Seed development was affected as well, as some seeds were quite small and did not germinate (Fig S1C).

To determine whether the mutation responsible for the SCN defect was attributable to a single gene or multiple genes, we crossed the mutant with wild type Col-0 and scored seedlings without GFP signals in the QC in the F2 segregation population. Intriguingly, among 404 F2 seedlings, only 19 seedlings could be reliably identified as mutant, resulting in a mutant frequency of about 1/20, which is dramatically less than the expected percentage for a single recessive mutation (1/4). A plausible explanation for the observed low percentage of mutants in the F2 population is that some mutants were lost because some mutant seeds could not germinate. Since phenotypic variation was also observed in the original mutant, albeit to a lesser extent, another explanation is that the mutation has incomplete penetrance; yet, another possibility is that there are other mutations that interact with the causal mutation, and these modifiers could alter the mutant/wildtype ratio in the F2 segregation population.

### 2. The gene with the causal mutation in our mutant is *TEN1/MDO1*

To distinguish among the possibilities described above and, more importantly, to determine whether the mutant phenotype can be attributed to a major allele, we performed an SSLP (Simple Sequence Length Polymorphism) analysis with an F2 segregation population resulting from a cross between the mutant and the wild type in the L*er* background. This analysis was first conducted with 44 mutant seedlings showing a clear SCN defect phenotype, using 14 markers that are located in different regions of each of the five chromosomes. Only markers F28J9A on chromosome one showed a recombinant rate significantly less than 50%, suggesting that the causal mutation was located on this chromosome (Fig 2). To corroborate this conclusion, we analyzed the same population of mutants using two additional SSLP markers, NGA128 and T2K10. The recombinant rate was 3.49% for T2K10, and 1.14% for NGA128. These results suggest that the No. 74 mutant phenotype is most likely due to a recessive mutation in a single gene located near the NGA128 marker (Fig 2).

**Fig 2.**
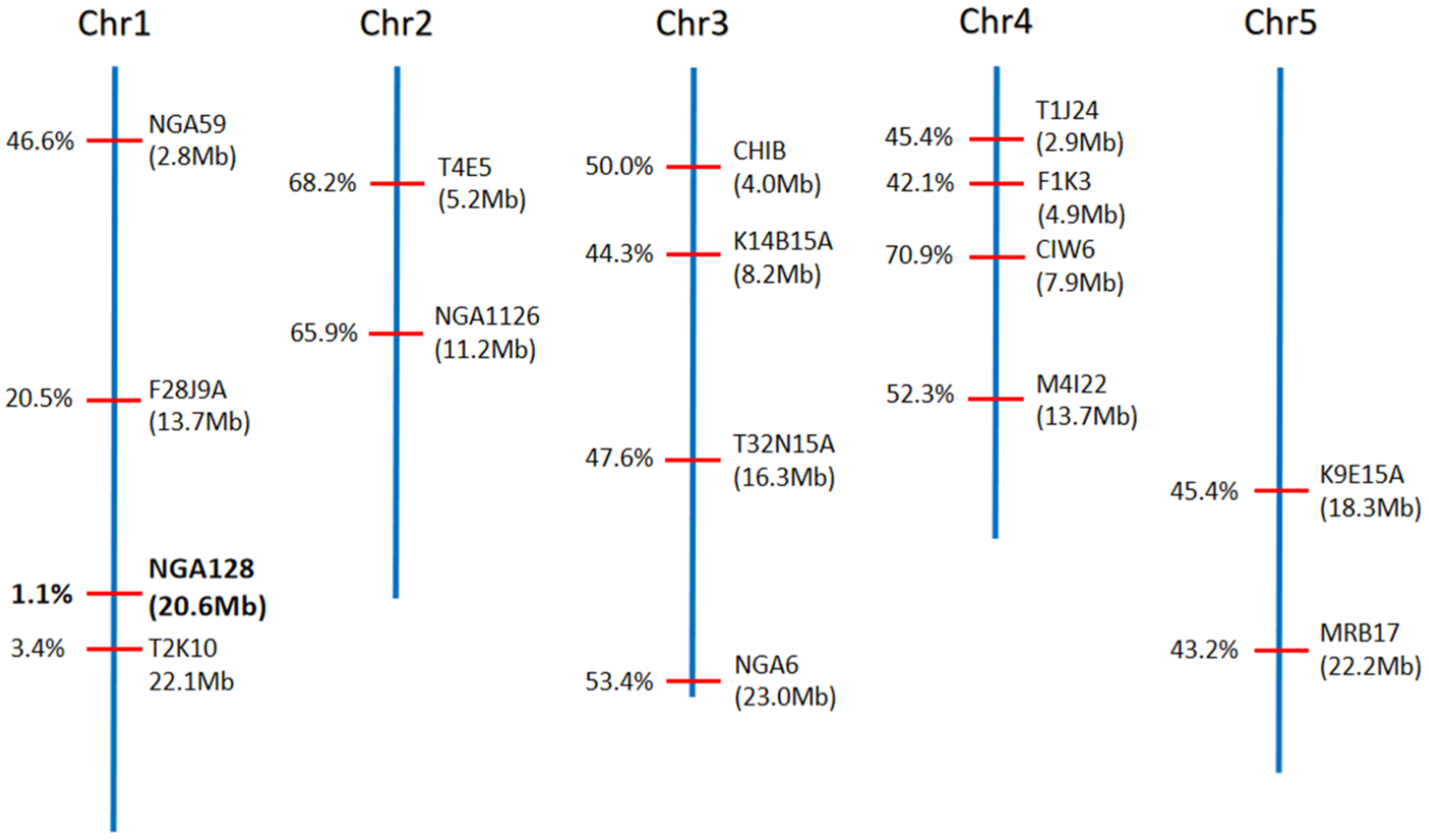
SSLP-assisted mapping located the causal mutation in mutant 74 near the NGA128 marker on chromosome one. The numbers on the left of each marker are the recombination frequency, and those below the markers (in the parenthesis) are their physical locations.

To pinpoint the location of the causal mutation, we next conducted whole genome resequencing with the mutants and their siblings from a F2 segregation population, as well as their parent lines. Bulked Segregant Analysis (see Methods for detail) clearly showed that the causal mutation is located in a region in chromosome one associated with NGA128 marker (Fig S2). This result is consistent with the genetic analysis described above, lending further support to the notion that the causal mutation in mutant 74 is within a single gene. To identify the gene with the causal mutation, we first searched the region defined by the BSA analysis for genes that harbor a G-to-A mutation that is characteristic of the EMS mutagenesis, and a SNP index value of one that indicates complete linkage between the mutated sites and causal mutation. Among the genes meeting these criteria, five genes were further investigated because they either contain nonsense mutations within their coding sequences (*AT1G53282, AT1G54030 (ERMO3*) and *AT1G54350 (ABCD2)*), or lost the start codon (*AT1G55720 (CAX6)*) (Table S2). *AT1G56260* (*TEN1*) caught our attention as it has been previously identified as *MERISTEM DISORGANIZATION 1* (*MDO1*), which plays an essential role in SCN maintenance (Hashimura and Ueguchi, 2011; Leehy et al., 2013). Remarkably, our mutant had the same G-to-A mutation in the 77^th^ codon as the *mdo1-1* mutant (Leehy et al., 2013), resulting in Gly to Glu substitution. Similarly to our mutant, the *mdo1-1* mutant affects SCR expression specifically in the SCN (Hashimura and Ueguchi, 2011). The *mdo1* mutant also resembles our mutant in many other respects, such as fascinated inflorescence stem and clustered flowers, as well as variable and incomplete penetrance of the mutant phenotypes (Hashimura and Ueguchi, 2011). We therefore postulated that the gene with the causal mutation in our mutant is very likely to be *MDO1*. Nevertheless, in complementation tests we have examined the function of all five candidate genes. In our first batch of transformations, however, we were unable to obtain any transgenic plants, which we believe is attributable to the low fertility of the mutant that was aggravated after agrobacteria infiltration. Therefore we next transformed an F2 segregation population of plants from a cross between the mutant and Col-0. Our reasoning is that, if the mutant is rescued, none of the resulting transgenic plants would display the mutant phenotype; otherwise, 1/20 would be abnormal, which is the percentage of mutants we have observed in an F2 segregating population. As shown in Table 1, only AT1G56260 was able to complement the mutant. Hence, we conclude that our mutant is the same as *mdo1-1*.

**Table 1.**
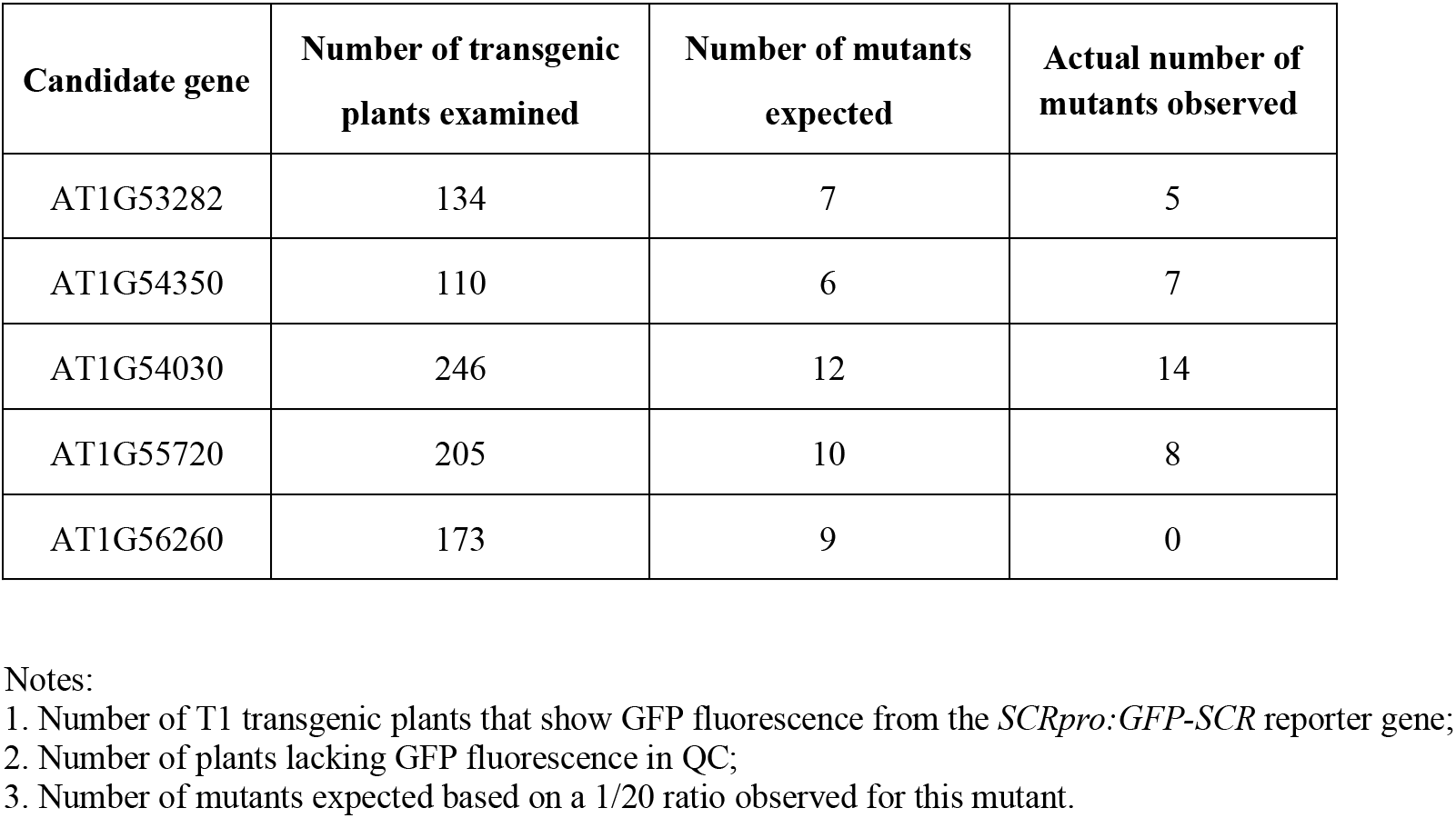
Summary of the complementation test results.

| Candidate gene | Number of transgenic plants examined | Number of mutants expected | Actual number of mutants observed |
| --- | --- | --- | --- |
| AT1G53282 | 134 | 7 | 5 |
| AT1G54350 | 110 | 6 | 7 |
| AT1G54030 | 246 | 12 | 14 |
| AT1G55720 | 205 | 10 | 8 |
| AT1G56260 | 173 | 9 | 0 |

Notes:
1. Number of T1 transgenic plants that show GFP fluorescence from the *SCRpro::GFP-SCR* reporter gene; 2. Number of plants lacking GFP fluorescence in QC; 3. Number of mutants expected based on a 1/20 ratio observed for this mutant.

### 3. TEN1/MDO1 has several conserved motifs essential to its function

TEN1 is a component of the so called CST complex that binds to and ensures the integrity of the telomere (Lim and Cech, 2021). The fact that our mutant has the same mutation (G77E) as the *mdo1* mutant underscores the critical importance of this amino acid residue to TEN1/MDO1 function. Indeed, there is evidence that the G-to-E mutation caused protein instability in TEN1 (Lim and Cech, 2021) and loss of TEN1’s ability to interact with CTC, a scaffold protein essential for the assembly of the CST complex (Lim and Cech, 2021).

Underscoring its essential role in telomere integrity, TEN1 is found in all organisms that have linear chromosomes (Prochazkova Schrumpfova et al., 2019). Bioinformatics analysis of *TEN1* homologs from a number of organisms, including yeast, plants and mammals, showed that G77 is located within a stretch of amino acid resides that are highly conserved in these organisms (Leehy et al., 2013). This motif was shown to be critical to TEN1 function, as the G77E mutation causes protein instability and loss of the ability to interact with STN1 (Leehy et al., 2013). The same analysis also revealed the presence of additional conserved motifs, but their functional importance has not been experimentally tested. To determine whether these additional motifs are indispensable for TEN1 function in plants, we identified the most conserved amino acid residues in them by multi-sequence alignment of *TEN1* homologs from a wide spectrum of plants, ranging from primitive land plants to seed plants, as well as two algae species and the fission yeast (Table S3). Two amino acid resides were found to be present in all TEN1 homologs: one is an arginine at position 21 (R21), the other is a glycine at position 100 (G100) (Fig 3A).

**Fig 3.**
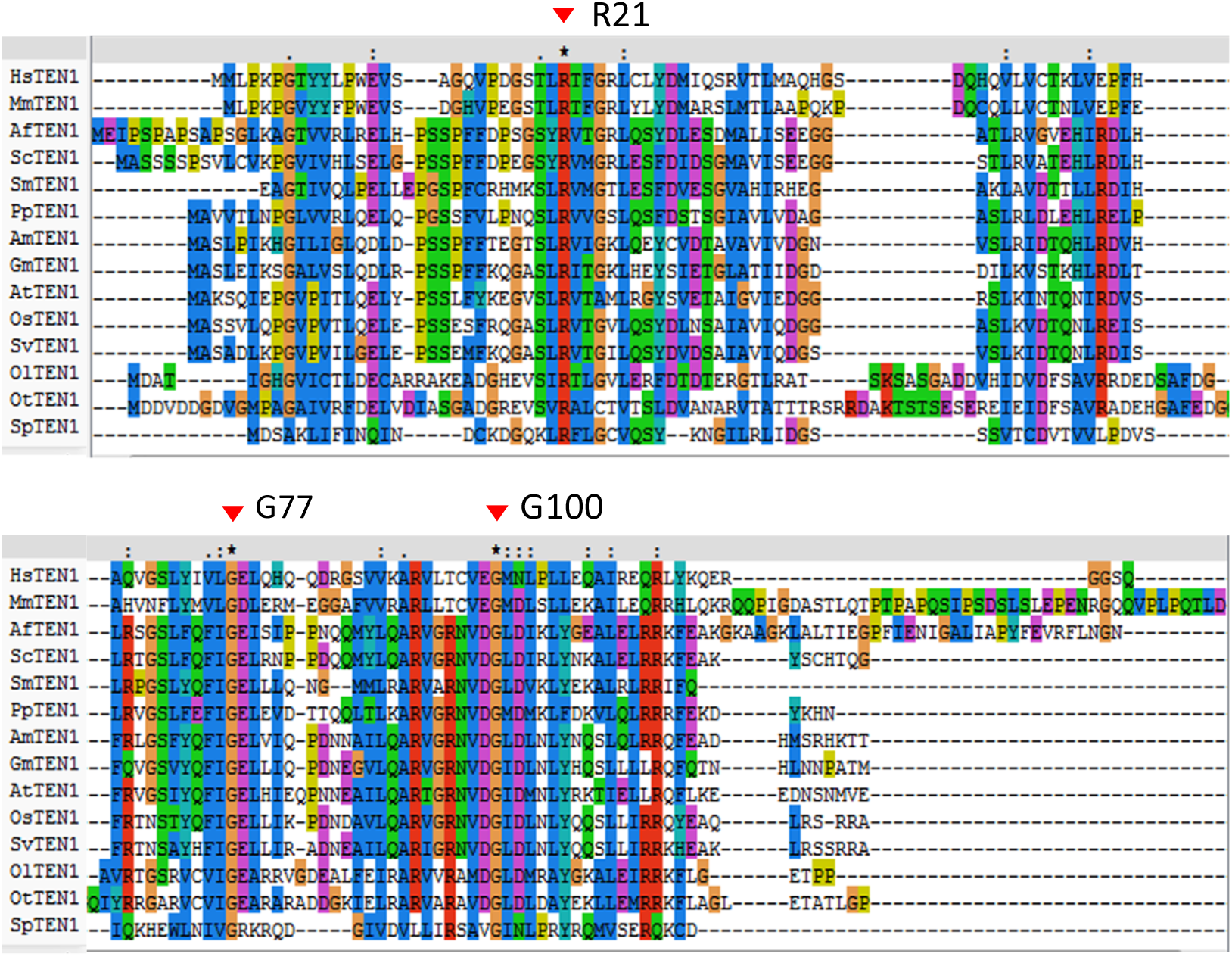
Multi-sequence alignment of TEN1 homologs from a diverse range of eukaryotes. Arrows mark the position of the most-conserved amino acids, R21 and G100, as well as G77, which is mutated in the *mdo1-1/ten1-3* mutant. See Table S3 for the full names of the species and their corresponding taxonomy groups.

To test the functionality of R21 and G100, we mutated the arginine to glycine (R21G) or glutamine (R21E), the glycine to glutamine (G100E), and then expressed the mutant forms as GFP-fusion proteins under the control of the *TEN1* promoter in the *TEN1* mutant. To circumvent the sterility issue of this mutant, we used the strategy described above (i.e., transforming F2 plants from a cross between the mutant and the wild type, and calculating the percentage of transgenic plants still showing mutant phenotypes). The results showed that all but the protein containing the G100E mutation were able to fully rescue the mutant (Table 2). Since the number of seedlings containing the transgene and showing mutant phenotypes was the same as that expected based on the 1/20 frequency that is characteristic of our mutant, we conclude that the G100E mutation has completely abolished the function of the TEN1 protein. This result suggests that G100 and hence the corresponding motif is critical for TEN1 function.

**Table 2.** Summary of results from the complementation tests for TEN1 variants.

| Vector | Number of transgenic plants examined | Number of mutants expected | Number of mutants observed |
| --- | --- | --- | --- |
| *TEN1pro::TEN1-eGFP* | 109 | 5 | 0 |
| *TEN1pro::R21G-eGFP* | 115 | 6 | 0 |
| *TEN1pro::R21E-eGFP* | 117 | 6 | 0 |
| *TEN1pro::G100E-eGFP* | 110 | 6 | 6 |

Notes:
1. Number of T1 transgenic plants that show GFP fluorescence from the *SCRpro::GFP-SCR* reporter gene; 2. Number of plants lacking GFP fluorescence in QC; 3. Number of mutants expected based on a 1/20 ratio observed for this mutant.

The TEN1 protein is mainly localized in the nucleus (Leehy et al., 2013). Hence, it is possible that the G100E mutation could cause a change in subcellular localization. To test this possibility, we examined by confocal microscopy the subcellular localization of the GFP fusion proteins, with or without the G100E mutation, in root epidermal cells of the transgenic plants used for the complementation tests (*TEN1pro::TEN1-eGFP (G100E)* and *TEN1pro::TEN1-eGFP*). As expected, the TEN1-GFP protein was found mainly in the nucleus, but it appeared to be also present in the cytoplasm and even in plasma membrane (Fig 4A). Strikingly, we found that the TEN1-GFP (G100E) protein was barely detectable, although it had the same subcellular localization as the wild type form (Fig 4, A and B). The low GFP fluorescence was unlikely due to the GFP tag, because the expression levels of the TEN1-GFP fusion proteins containing R21G (*TEN1pro::TEN1-eGFP* (*R21G*)) or R21E (*TEN1pro::TEN1-eGFP* (*R21E*)) were similar to that of TEN1-GFP ((Fig 4A).

**Fig 4.**
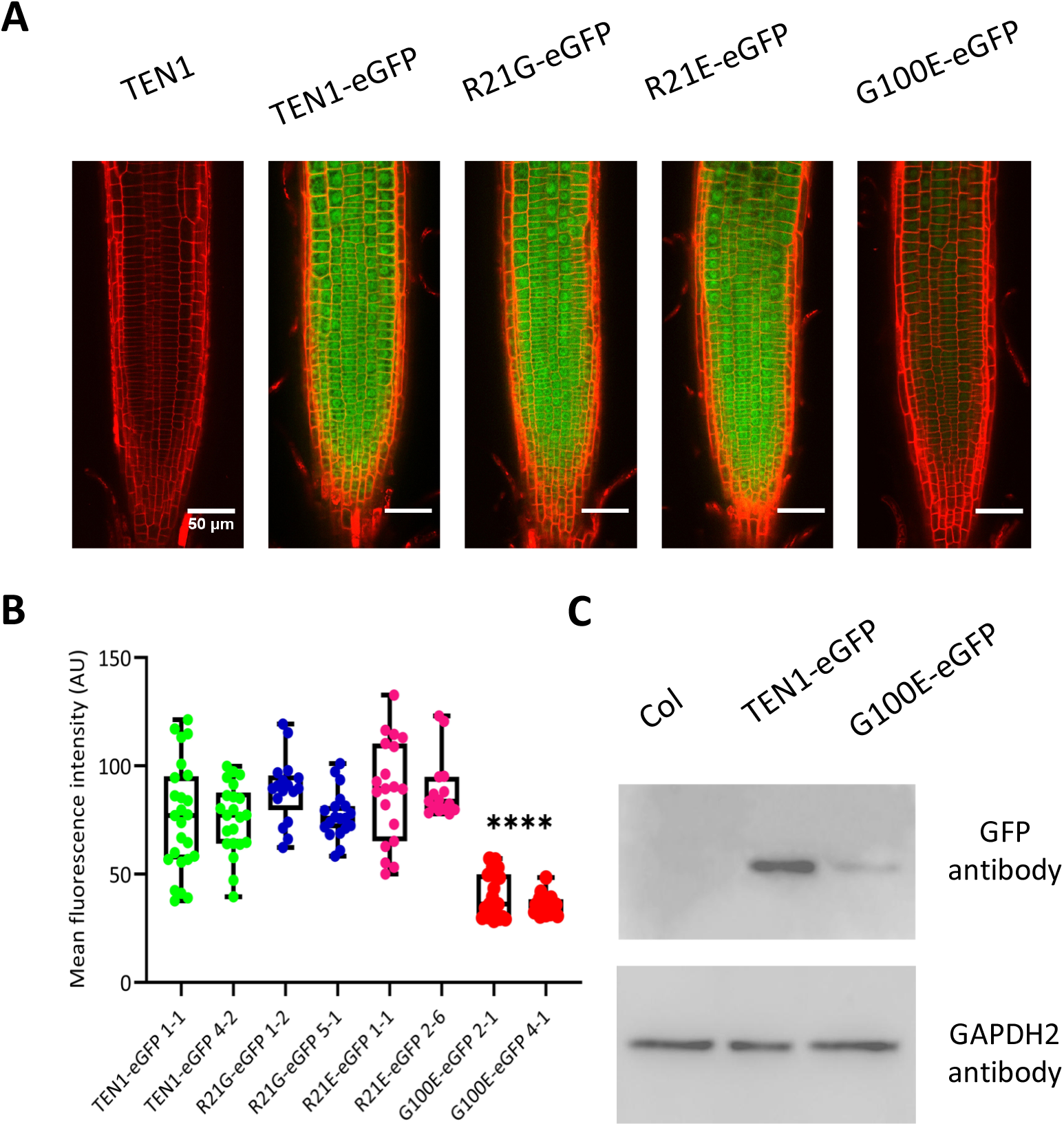
G100 is essential to TEN1 protein expression. A. Representative confocal microscope images showing the expression of the different GFP fusion proteins. Bar = 50 μm. B. Box plot and statistical analysis of GFP signal intensity in the root tip of transgenic plants. AU, artificial unit. \*\*\*\**p* < 0.0001. *t*-test. n ≥ 15. C. Western blot assay showing the protein level of the TEN1-GFP fusion protein with the G100E mutation (G100E-eGFP), relative to non-transgenic plants (Col), and the wild type TEN1-eGFP fusion protein. GAPDH2 was used as a loading control

The observed low fluorescence in the TEN1-eGFP (G100E) transgenic plants could be attributed to a low level of protein or to misfolding of the protein. To distinguish between these possibilities, we compared the protein level of the TEN1-GFP proteins with or without the G100E mutation by Western blot. As shown in Fig 4C, the mutated protein had a much lower level relative to the wild type GFP fusion protein. Since the G77E mutation is known to affect the physical interaction between TEN1 and STN1 (Leehy et al., 2013), we reasoned that the G100E mutation may have a similar effect, which would make the protein more vulnerable to degradation. Surprisingly, however, yeast two-hybrid assay indicated that the G100E mutant form was still able to interact with STN1 and the interaction was as strong as that between the wild type form and STN1 (Fig 5A). To validate this result, we tested their interaction further by bimolecular fluorescence complementation. As shown in Fig 5B, the G100E mutation did not affect the interaction between TEN1 and STN1. These results suggest that the low level of the G100E mutant form is not due to its dissociation from the CST complex. Consistent with the result from our confocal microscope observation (Fig 4A), TEN1-GFP was clearly detected in the plasma membrane and cytoplasm in addition to its nuclear localization (Fig 5B).

**Fig 5.**
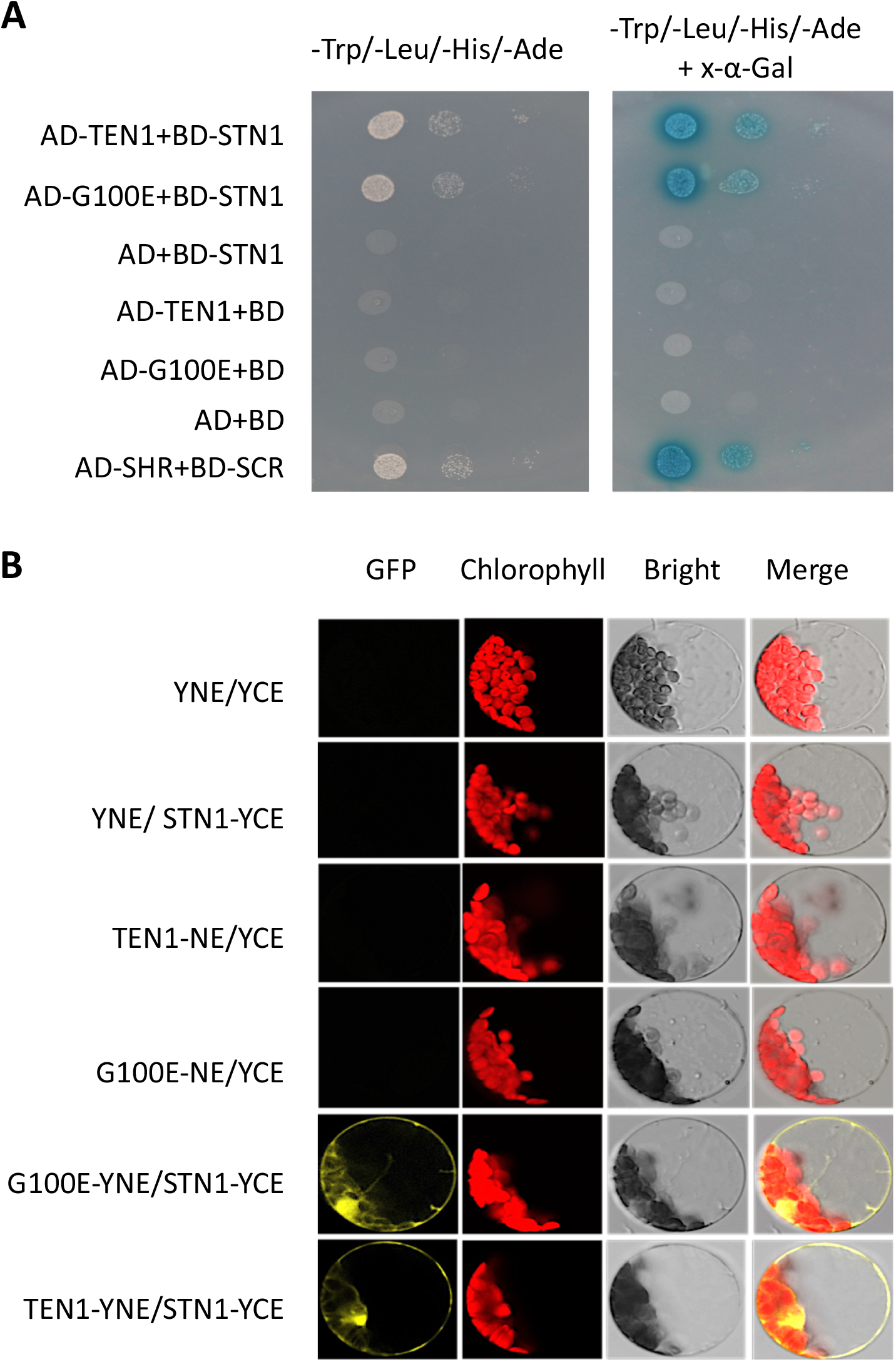
G100E mutation does not affect physical interaction between TEN1 and STN1. A. Yeast two-hybrid assay showing that the G100E mutation does not affect the ability of TEN1 to interact with STN1. The SHR and SCR pair serves as the positive control. YNE, N-termus of YFP; YCE, C-terminus of YFP. B. Bimolecular fluorescence complementation assay for the interaction between STN1 and TEN1 with or without the G100E mutation.

The steady-state level of TEN1 protein could be regulated by the ubiquitin–proteasome pathway and the G100E mutation may make it a better target for degradation. To test this hypothesis, we treated seedlings containing the *TEN1pro::TEN1-eGFP* or *TEN1pro:TEN1(G100E)-eGFP* construct with the proteasome inhibitor MG132 and then examined GFP fluorescence by confocal microscopy. As negative controls, the R21G and R21E mutation forms were also included in this experiment. No apparent difference in fluorescence signal intensity was detected for any of these proteins before and after MG132 treatments (Fig S3). These results suggest that TEN1 is degraded by a mechanism unrelated to the ubiquitin–proteasome pathway.

### 4. Other telomere protecting factors also play a role in stem cell maintenance in the root

The finding that mutation in TEN1 causes SCN defect raises the question of whether other telomere integrity factors also have a role in stem-cell maintenance in the root. To address this question, we examined the SCN in mutants for the other components of the CST complex, CTC and STN1, as well as telomerase. Compared to the wild type, all mutants displayed a short and variable root phenotype (Fig6, A and B). Confocal microscopic imaging showed that in *ctc1* and *stn1* mutants the STN was severely disorganized with cells heavily stained by propidium dye, indicating cell damage (Fig 6C). Cell damage was also observed in two telomerase mutants, *tert1* and *tert-2*, although their SCN appear to be morphologically normal. These results suggest that telomere integrity is essential to stem cell maintenance.

**Fig 6.**
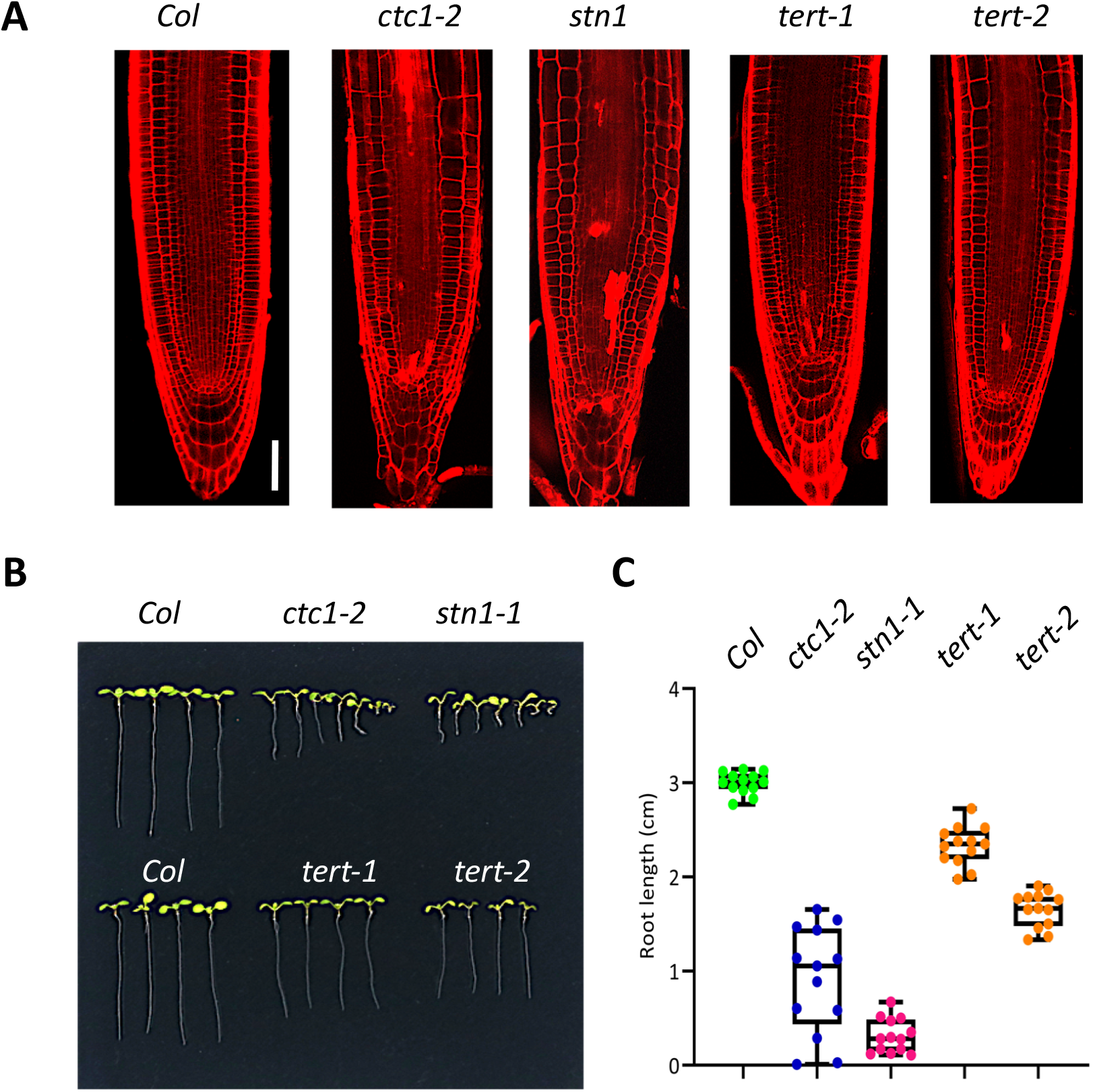
CTC1, STN1 and telomerase also play a role in SCN maintenance. A. Confocal microscope images showing disorganized SCN in *ctc1, stn1* and telomerase (*tert*) mutants. B. *ctc1*, *stn1* and two *tert* mutants have short roots. The seedlings were eight days old grown in MS medium. C. Box plots of root length of the root length of *ctc1*, *stn1* and two *tert* mutants. n=13

### 5. SCR maintains the SCN by maintaining telomere integrity

The finding that mutation in TEN1 causes SCN defect raises the possibility that SCR may maintain the SCN by maintaining the expression level of TEN1 and other telomere protecting factors. To investigate this possibility, we first examined *TEN1* expression in the *scr* mutant by qRT-PCR. Indeed, we found that *TEN1* transcript level was dramatically reduced in the *scr-1* mutant (Fig 7A). We also examined the expression pattern and level of TEN1 in the *scr* mutant using the *TEN1pro::TEN1-GFP* reporter construct. Consistent with the qRT-PCR result, we could barely detect any GFP signal in the SCN region in *scr* root (Fig 7B).

**Fig 7.**
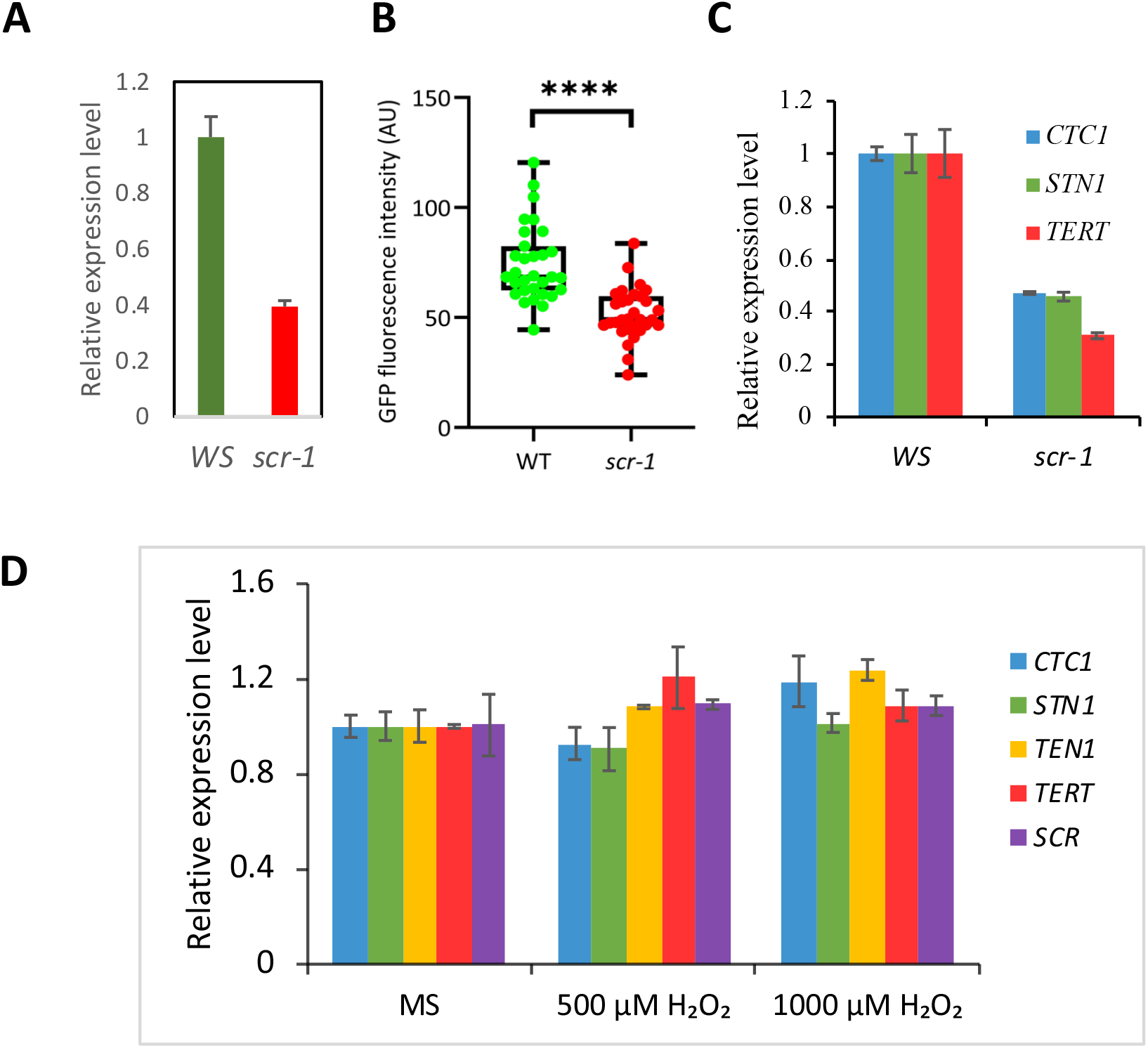
*TEN1, CTC1, STN1* and telomerase are downregulated in the *scr* mutant. A. qRT-PCR assay of *TEN1* transcripts in the roots of wild type (Ws) and *scr* seedlings. B. TEN1-eGFP protein level in wild type (WT) and *scr*, as determined by the intensity of green fluorescence. C. Transcript levels of *CTC1, STN1* and telomerase (*TERT*) in the roots of wild type and *scr* seedlings. D. qRT-PCR of *TEN1, CTC1, STN1* and telomerase in the roots of wild type seedlings with or without H_2_O_2_ treatments. For qRT-PCR, *actin 7* was used as an internal control.

The SCN defect in the *scr* mutant is unlikely attributable only to the lower expression of TEN1, because *scr* has a much shorter root than *ten1*. We therefore also examined the transcript levels of STN1 and CTC1 as well as telomerase. As expected, all of these genes were found to be dramatically downregulated in *scr* (Fig 7C). These results suggest that SCR maintains the SCN by maintaining the integrity of chromosome ends.

Recently we showed that the *scr* mutant has a defect in redox homeostasis, which partly explains its shorter root phenotype (Fu et al., 2021). It is thus possible that the reduction in expression of telomerase and CST complex is an indirect consequence of an elevated level of reactive oxygen species in the mutant. To test this, we compared the transcript level of telomerase, *TEN1*, *STN1* and *CTC1* in wild-type seedlings grown in MS medium or H_2_O_2_-containing medium. As shown in Fig 7D, none of these genes was affected transcriptionally by H_2_O_2_ at a concentrations of 500 uM and 1000 uM, which have apparent growth inhibitory effects but are still below the level causing cell death (Cui et al., 2014). This result lends support to the notion that SCR maintains the SCN by maintaining telomere integrity.

If the reduction in TEN1 expression is a cause for the SCN defect in the *scr* mutant, the *scr* mutant could have an elevated level of damaged DNA and thus becomes hypersensitive to conditions that elicit DNA damage. To see whether this is the case, we treated *scr* mutant and wild-type (Col) seedlings with zeocin, a reagent known to cause DNA damage. As shown in Fig 8, *scr* seedlings became yellow at 20 ug/mL of zeocin and bleached at 50 ug/mL, whereas the wild type had no lesions. This result lends support to the notion that the *scr* mutant indeed has a defect in genome stability. It further suggests that reduced expression of telomerase and the CST complex account, at least partially, for the SCN defect in the *scr* mutant.

**Fig 8.**
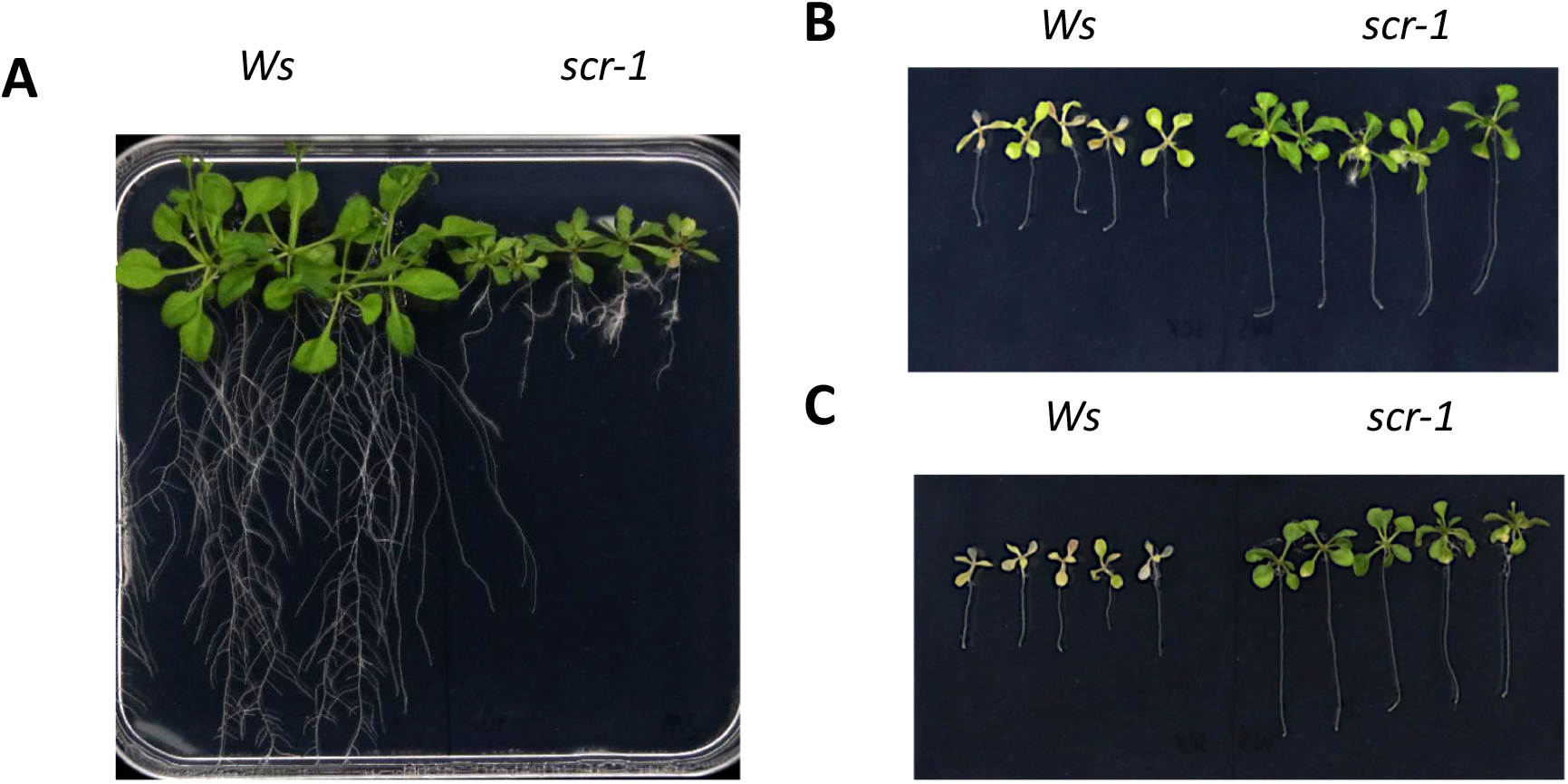
The *scr* mutant is hypersensitive to zeocin, a DNA damage reagent. A. Wild type (Col) and *scr* seedlings 22 days after germination in MS medium. B and C. Wild type (Col) and *scr* seedlings 14 days after 8-days-old seedlings were transferred to MS medium containing 20 (B) or 50 (C) μg/mL zeocin.

## Discussion

Maintaining the SCN is essential to postembryonic growth and development in plants. Although SCR was identified as a key regulator of stem cell maintenance in *Arabidopsis* root more than two decades ago, the mechanism by which SCR maintains the root SCN has remained unclear. Serendipitously, in this study we discovered a connection between SCR and telomere protecting factors. In a genetic screen aimed to identify factors that regulate SCR expression, we uncovered a mutant that has lost SCR expression in the SCN, resulting in disorganized SCN and a short root phenotype. Through subsequent molecular analyses we were able to locate the causal mutation within *TEN1* that encodes a component of the telomere-end protecting CST complex. Because of the similar SCN defects in *ten1* and *scr* mutants, we wondered if altered TEN1 expression and other telomere protecting factors could be a cause of the SCN defect in *scr*. We tested and corroborated this hypothesis by RT-PCR, which showed that telomerase and the three components of the CST complex, TEN1, CTC1 and STN1, were all downregulated in *scr*. This change in gene expression is clearly not due to an elevated level of reactive oxygen species that accumulated in the *scr* mutant (Fu et al., 2021), because none of these genes was induced by hydrogen peroxide. These results together suggest that SCR maintains the SCN, at least partly, by sustaining optimal expression level of telomere protecting factors and thus ensuring telomere integrity. Supporting this conclusion, we demonstrated that the *scr* mutant was hypersensitive to Zeocin, a DNA damage reagent, which would be expected if telomere integrity is compromised.

The role of SCR in maintaining genome stability may not be limited to the telomere because one of its downstream targets, STN1, has been shown to interact with DNA polymerase alpha, an enzyme involved in genome replication (Derboven et al., 2014). In addition, there is evidence that individual components of the CST complex may have distinct functions that are currently not understood. For instance, protein subcellular location studies showed that in the nucleus TEN1 and CTC1 are enriched in many spots beyond the telomeres, which do not overlap (Leehy et al., 2013; Surovtseva et al., 2009). In the present study we found that TEN1 protein was even localized in the plasma membrane and cytoplasm (Fig 4A and E). These observations suggest that TEN1 is a multifunctional protein, which was also proposed in a recent study by others (Lee et al., 2016). Considering the fact that TEN1, CTC1 and STN1 are downregulated in *scr*, it is logical to postulate that SCR maintains the root SCN by ensuring genome stability at the genome scale.

In this study we also identified and functionally validated the indispensability of a conserved motif in TEN1. Through phylogenetic analyses of TEN1 homologs from a wide range of eukaryotes, we showed that three amino acid resides are present in all plants: R21, G77 and G100. The functional importance of G77 is underscored by the fact that it is exactly the same mutation in our mutant and the *mdo1* mutant (Leehy et al., 2013). To examine the functionality of R21 and G77, we substituted glutamic acid (E) or Alanine (A) using site-directed mutagenesis and expressed them in our mutant. Only the G100E mutation failed to complement the *ten1* mutant, indicating that G100 is critical for TEN1 function. Interestingly, in further experiments we showed that the TEN1 protein with the G100E mutation was barely detectable, although this mutation did not affect the physical interaction between TEN1 and STN1. These results strongly suggest that the motif containing G100 is required for protein stability, although how G100 exerts this role is still unknown. It is noteworthy that components in the CST complex interact not only with each other but also with other proteins, such POT1 (Renfrew et al., 2014) and DNA polymerase (Derboven et al., 2014). It is thus possible that this new motif is required for protein-protein interaction with other proteins, which warrants further investigation.

## Materials and Methods

### 1. Plant growth conditions and treatments

For root growth experiments, seedlings were grown aseptically in Murashige and Skoog (MS) medium supplemented with 1% sucrose and 0.6% Phytagel (Beijing BioDee Biotech, P8169) in square petri dishes, which were placed vertically in a Percival growth chamber (model 41L). The growth conditions were 16-h light (50 micromoles/ m^2^/ sec of light irradiance) and 8-h darkness with a constant temperature of 22 °C. For this purpose, seeds were first sterilized with 10% bleach, then washed thoroughly with sterile H_2_O before sowing. For bulking and experiments with above-ground organs, plants were grown in a growth room with the same settings as the growth chamber.

For chemical treatments, seedlings were first germinated and grown in 1x MS medium for 6 days (for H_2_O_2_) or 8 days (for zeocin and MG-132, a peptide-aldehyde proteasome inhibitor). After transfer to the appropriate medium, seedlings were then allowed to growth for three hours, four or six days with MG-132 (MedChemExpress, HY-13259), H_2_O_2_ or Zeocin (Coolaber, SL4140), respectively. As a control, a similar number of seedlings were also transferred to fresh MS medium and mock treated for the same amount of time.

### 2. EMS mutagenesis, mutant screen and characterization

Ethyl methanesulfonate (EMS) mutagenesis was conducted according to the Arabidopsis handbook (Weigel and Glazebrook, 2002). Approximately 20,000 seeds homozygous for the *SCRpro::GFP-SCR* transgene were treated overnight at room temperature with 10 mL 0.1% ethyl methanesulfonate (EMS, Sigma M0880, MO, USA). After thorough washing, the seeds were suspended in 0.01% agarose and sowed in soil (three to five seeds per pot). At maturity, the seeds (M0) from all the plants in each pot were pooled and numbered. For mutant screening, the root tip (~1 cm) of one-week-old seedings grown in MS medium was cut and examined under a compound fluorescence microscope (Olympus BX61), and seedlings showing abnormal pattern or intensity of GFP fluorescence (M1) were transferred to soil for seed setting. Putative mutants were re-examined at the M2 generation and true mutants were crossed to L*er* for subsequent genetic analyses and causal gene identification. For confocal microscopy, seedlings were stained for one minute with propidium iodide (Sigma, P-4170) dissolved in ddH_2_O at a concentration of 10 μg/ml and images were captured using a Leica SP8 confocal microscope.

### 3. Genetic analysis and marker-assisted mapping of the causal genes in mutant 74

For this purpose, mutants in the F2 population from a cross between the mutant 74 (in the Col background) and L*er* were selected based on the expression pattern of GFP fluorescence in the root tip. The percentage of seedlings showing loss of GFP-SCR expression was then calculated as a proxy for the penetrance of the mutant phenotype. To determine the chromosomal location, individual mutants were genotyped when they were one month old by PCR using SSLP markers. Primer information was retrieved from the Arabidopsis Information Resource (TAIR, http://www.arabidopsis.org) and is listed in Table S1. DNA was extracted using the CTAB method.

### 4. High-throughput sequencing and BSA analysis

Three groups of plants were sequenced using the Illumina paired-end sequencing method on the HiSeq4000 platform: the same F2 mutant plants as described above for marker-assisted mapping (57 plants), normal F2 plants (30 plants), and the original mutant 74 (M2, 30 plants). An equal amount of leaf samples was collected from each plant, and the leaf samples were pooled for each group for DNA extraction using the TIANGEN DNA purification kit (TIANGEN, DP305-02). High-throughput sequencing and Bulked Sequence Analysis (BSA) were provided by a commercial service (Lianchuan Biotech Ltd., Hangzhou, China).

After removal of linker sequences, contamination and low-quality reads, 4.28 Gbp, 3.44 Gbp and 2.12 Gbp valid sequence reads were obtained for the three groups of samples, respectively, corresponding to a coverage of 56.5X, 45.3X and 28.0X. After the sequence reads were mapped to the Arabidopsis genome usingTAIR10 (https://www.arabidopsis.org/download/), SNPs in the mutants were then identified by pairwise comparison, for which the genome sequence of L*er* that we generated recently was also used as the reference (Li et al., 2020). For a particular mutated site, the ratio between the number of reads with that nucleotide and the total number of reads was calculated as the SNP index. Thus, the SNP_index should be one for those sites that are completely linked to the causal mutation, and 0.33 for the normal F2 population. The Delta_SNP_index was then calculated by subtracting the SNP_index for F2 normal form that for F2 mutants. Finally, SNPs with a Delta_SNP_index within the top 0.5% that also causes missense or nonsense mutation were selected as candidate causal mutations and the genes containing these SNPs were subject to further investigation.

### 5. Complementation tests

All genes tested in this study were expressed in mutant 74 under the control of their own promoters. For the promoter, the intergenic region up to 3kb, upstream of the first codon, was PCR amplified. For *AT1G54030, AT1G55720* and *AT1G56260*, the coding region was amplified from genomic DNA, whereas for *AT1G54350* the cDNA was used as the template. *AT1G53282* is an exception because of its small size: its promoter and coding region were amplified as a single piece using genomic DNA as the template. For the convenience of cloning, appropriate restriction sites were introduced into the primers used for the PCR amplification (Table S1). After digestion with corresponding enzymes and purification, PCR fragments were cloned into the pBluescript vector. Finally, clones whose sequences have been confirmed by Sanger sequencing were subcloned into the expression vector pCambia1305 or pCambia1302 (see Table S1 for the restriction sites used).

The construct for expressing TEN1-GFP fusion protein was generated in a similar manner to the clones above, except that the coding region was PCR amplified using the forward primer TEN1-F and a reverse primer that does not contain the stop codon (TEN1-R, Table S1). The GFP sequence in the expression vector was also retained and the two sequences were fused in frame. To construct GFP tagged mutant versions of TEN1, the point mutation was introduced by overlapping PCR. First, two fragments were separately amplified: a 5’ fragment using TEN1-F and a reverse primer that contains the mutant nucleotide (Rm primers), and a 3’ fragment using TEN1-R and a forward primer that contains the mutant nucleotide (Fm primers); The two PCR reactions were then mixed and subjected to PCR for five cycles; Finally, fresh TEN1-F and TEN1-R primers were added to this mix and PCR was allowed to run for 28 cycles.

For cloning described in this study, a high-fidelity DNA polymerase, PrimeSTAR®HS DNA Polymerase (Takara, R010A), was used for all PCR reactions. Transgenic plants were generated in the Col–0 ecotype using the flower dip method (Clough and Bent, 1998) and selected on MS medium containing kanamycin at a concentration of 50 μg/mL^−1^.

### 6. Other molecular assays

#### 6.1. RNA extraction and quantitative RT-PCR

Total RNA was extracted from the roots of 8-day seedlings using the Trizol reagent. After treatment with DNase I (Thermo Scientific, EN0521) to remove residual genomic DNA, the RNA was converted into cDNA using PrimeScript™ II 1st Strand cDNA Synthesis Kit (TAKARA, 6210A) following the instruction in the manual. The cDNA was then used as a template in subsequent real time PCR assay, for which Taq Pro Universal SYBR qPCR Master Mix (Vazyme, Q711-02) was used on an Bio-Rad CFX Connect real-time system. The primers used in the assay were CTC1-RT-F and CTC1-RT-R for CTC1, STN1-RT-F and STN1-RT-R for STN1, TEN1-RT-F and TEN1-RT-R for TEN1, TERT-RT-F and TERT-RT-R for TERT, and SCR-RT-F and SCR-RT-R for SCR. 18S rRNA was used as an internal control for most qRT-PCR reactions, except for the H_2_O_2_, where *ACTIN7* was used as the control. See Table S1 for information about the primers.

#### 6.2 Yeast two-hybrid assay

The MATCHMAKER two-hybrid system 3 (http://www.bdbiosciences.com) was used for this experiment, whereby STN1 was used as the bait and TEN1 with or without the G100E mutation was used as the prey. The STN1 and TEN1 coding sequences were amplified by PCR using cDNA made from wild type Col seedlings as the template, cut with EcoRI along with PstI or BamHI respectively, and then cloned into the corresponding sites of the GBKT7 or GADT7 vectors. After confirmation by Sanger sequencing, the STN1-BD and TEN1-AD or TEN1-G100E-AD plasmids, as well as negative controls, were co-transformed into competent yeast cells (strain: YH109), and cells with both constructs were selected on SD/-Leu/-Trp medium (Clontech, 630417). Ten colonies picked up with a tooth pick were diluted in 1ml 0.9%NaCl and spotted onto SD/-Ade/-His/-Leu/-Trp medium (Clontech, 630428) with or without a-X-Gal (GoldBio, 107021-38-5). See Table S1 for information about the primers.

#### 6.3 Bimolecular fluorescence complementation

This assay was conducted essentially as described (Walter et al., 2004). TEN1 with or without the G100E mutation was fused to the N-terminal 155 amino acids of YFP, which were named as TEN1-YNE and G100E-YNE respectively, whereas STN1 was fused to the C-terminal 86 amino acids of YFP, and the fusion protein was named as STN1-YCE. To make the TEN1-YNE or G100E-YNE expressing construct, TEN1 with or without the G100E mutation was PCR amplified using primers TEN1-BiFC-F and TEN1-BiFC-R (Table S1) and their clones described above as the templates, cut with BamHI and XhoI that had been introduced into the primers during primer synthesis, and cloned into the same restriction sites in the pSPYNE-35S vector. The STN1-YCE expressing construct was cloned into the pSPYCE-35S vector by introducing XbaI and XhoI into the coding region of STN1 by PCR using the STN1-BiFC-F and STN1-BiFC-R primer pair (Table S1). The plasmids for STN1-YCE and TEN1-YNE or G100E-TNE were then introduced together into Arabidopsis protoplasts prepared according to (Yoo et al., 2007), and YFP fluorescence as well as chlorophyll autofluorescence were then imaged using a Leica TCS SP8 confocal microscope. As controls, a number of plasmid pairs were also co-transformed into the protoplast, including the YNE+YCE pair, TEN1-YNE+YCE, G100E-YNE+YCE, and YNE+STN1-YCE pairs.

#### 6.4. Western blot

Eight-day old seedlings grown in MS medium were used for this experiment. For each sample, 0.1 g of root cut 1cm from the tip was grounded in 300 μL ice-cold protein extraction buffer (50mM Tris-HCl, pH7.5, 150 mM NaCl, 1% Triton X-100, 1 mM EDTA, 1X protease inhibitor cocktail, and 1 mM PMSF). After centrifugation (12,000 rpm, 4°C, 10 min.), the supernatant was mixed with SDS loading buffer, boiled for 5 min., and then loaded to 10% SDS-polyacrylamide gel for electrophoresis. The proteins were then transferred for 1 h to a PVDF membrane (MilliporeSigma™ ISEQ00010) using the semi-dry method, and the GFP fusion proteins were detected using the JL-8 (Clontech, 632380; 1:2500 dilution) as the primary antibody and an HRP-conjugated Affinipure Goat Anti-Mouse antibody (Proteintech, SA00001-1; 1:2500 dilution) as the secondary antibody. As a loading control, GAPDH was also detected using an antibody against it from Proteintech (60004-1-Ig; 1:20000 dilution). Signal was visualized using the ECL Western Blotting Reagent (Hyyan Biotech, HY005) and a chemifluorescence imaging system from Syngene (Gbox Chemi XRQ).

## Acknowledgements

The authors are thankful to Jen Kennedy (Florida State University) for editing the manuscript. This research is supported by the National Science Foundation of China (grant no. 31871493), Florida State University, and Northwest Agriculture and Forest University.

## Notes

Funding: This research is supported by the National Science Foundation of China (grant No. 31871493), Florida State University, and Northwest Agriculture and Forest University.

## References

Aida, M., Beis, D., Heidstra, R., Willemsen, V., Blilou, I., Galinha, C., Nussaume, L., Noh, Y.S., Amasino, R., and Scheres, B. (2004). The PLETHORA genes mediate patterning of the Arabidopsis root stem cell niche. Cell 119, 109–120.

Benfey, P.N., and Scheres, B. (2000). Root development. Curr Biol 10, R813–815.

Clough, S.J., and Bent, A.F. (1998). Floral dip: a simplified method for Agrobacterium-mediated transformation of *Arabidopsis thaliana*. Plant J 16, 735–743.

Cui, H., Kong, D., Wei, P., Hao, Y., Torii, K.U., Lee, J.S., and Li, J. (2014). SPINDLY, ERECTA, and Its Ligand STOMAGEN Have a Role in Redox-Mediated Cortex Proliferation in the Arabidopsis Root. Mol Plant 7, 1727–1739.

Cui, H., Levesque, M.P., Vernoux, T., Jung, J.W., Paquette, A.J., Gallagher, K.L., Wang, J.Y., Blilou, I., Scheres, B., and Benfey, P.N. (2007). An evolutionarily conserved mechanism delimiting SHR movement defines a single layer of endodermis in plants. Science 316, 421–425.

Derboven, E., Ekker, H., Kusenda, B., Bulankova, P., and Riha, K. (2014). Role of STN1 and DNA polymerase alpha in telomere stability and genome-wide replication in Arabidopsis. PLoS Genet 10, e1004682.

Di Laurenzio, L., Wysocka-Diller, J., Malamy, J.E., Pysh, L., Helariutta, Y., Freshour, G., Hahn, M.G., Feldmann, K.A., and Benfey, P.N. (1996). The SCARECROW gene regulates an asymmetric cell division that is essential for generating the radial organization of the Arabidopsis root. Cell 86, 423–433.

Dolan, L., Janmaat, K., Willemsen, V., Linstead, P., Poethig, S., Roberts, K., and Scheres, B. (1993). Cellular organisation of the Arabidopsis thaliana root. Development 119, 71–84.

Fu, J., Zhang, X., Liu, J., Gao, X., Bai, J., Hao, Y., and Cui, H. (2021). A mechanism coordinating root elongation, endodermal differentiation, redox homeostasis and stress response. Plant J 107, 1029–1039.

Hashimura, Y., and Ueguchi, C. (2011). The Arabidopsis MERISTEM DISORGANIZATION 1 gene is required for the maintenance of stem cells through the reduction of DNA damage. Plant J 68, 657–669.

Helariutta, Y., Fukaki, H., Wysocka-Diller, J., Nakajima, K., Jung, J., Sena, G., Hauser, M.T., and Benfey, P.N. (2000). The SHORT-ROOT gene controls radial patterning of the Arabidopsis root through radial signaling. Cell 101, 555–567.

Kobayashi, A., Miura, S., and Kozaki, A. (2017). INDETERMINATE DOMAIN PROTEIN binding sequences in the 5’-untranslated region and promoter of the SCARECROW gene play crucial and distinct roles in regulating SCARECROW expression in roots and leaves. Plant Mol Biol 94, 1–13.

Lee, J.R., Xie, X., Yang, K., Zhang, J., Lee, S.Y., and Shippen, D.E. (2016). Dynamic Interactions of Arabidopsis TEN1: Stabilizing Telomeres in Response to Heat Stress. Plant Cell 28, 2212–2224.

Leehy, K.A., Lee, J.R., Song, X., Renfrew, K.B., and Shippen, D.E. (2013). MERISTEM DISORGANIZATION1 encodes TEN1, an essential telomere protein that modulates telomerase processivity in Arabidopsis. Plant Cell 25, 1343–1354.

Li, J., Wang, B., Zhu, X., Li, R., Fu, J., and Cui, H. (2020). Novel Regulators of Sugar-Mediated Lateral Root Development in *Arabidopsis thaliana*. Genes 11, 10.3390/genes11111257.

Lim, C.J., and Cech, T.R. (2021). Shaping human telomeres: from shelterin and CST complexes to telomeric chromatin organization. Nat Rev Mol Cell Biol 22, 283–298.

Long, Y., Goedhart, J., Schneijderberg, M., Terpstra, I., Shimotohno, A., Bouchet, B.P., Akhmanova, A., Gadella, T.W., Jr., Heidstra, R., Scheres, B., et al. (2015a). SCARECROW-LIKE23 and SCARECROW jointly specify endodermal cell fate but distinctly control SHORT-ROOT movement. Plant J 84, 773–784.

Long, Y., Smet, W., Cruz-Ramirez, A., Castelijns, B., de Jonge, W., Mahonen, A.P., Bouchet, B.P., Perez, G.S., Akhmanova, A., Scheres, B., et al. (2015b). Arabidopsis BIRD Zinc Finger Proteins Jointly Stabilize Tissue Boundaries by Confining the Cell Fate Regulator SHORT-ROOT and Contributing to Fate Specification. Plant Cell 27, 1185–1199.

Moreno-Risueno, M.A., Sozzani, R., Yardimci, G.G., Petricka, J.J., Vernoux, T., Blilou, I., Alonso, J., Winter, C.M., Ohler, U., Scheres, B., et al. (2015). Transcriptional control of tissue formation throughout root development. Science 350, 426–430.

Moubayidin, L., Di Mambro, R., Sozzani, R., Pacifici, E., Salvi, E., Terpstra, I., Bao, D., van Dijken, A., Dello Ioio, R., Perilli, S., et al. (2013). Spatial coordination between stem cell activity and cell differentiation in the root meristem. Dev Cell 26, 405–415.

Nakajima, K., Sena, G., Nawy, T., and Benfey, P.N. (2001). Intercellular movement of the putative transcription factor SHR in root patterning. Nature 413, 307–311.

Prochazkova Schrumpfova, P., Fojtova, M., and Fajkus, J. (2019). Telomeres in Plants and Humans: Not So Different, Not So Similar. Cells 8, 58.

Renfrew, K.B., Song, X., Lee, J.R., Arora, A., and Shippen, D.E. (2014). POT1a and components of CST engage telomerase and regulate its activity in Arabidopsis. PLoS Genet 10, e1004738.

Sabatini, S., Heidstra, R., Wildwater, M., and Scheres, B. (2003). SCARECROW is involved in positioning the stem cell niche in the Arabidopsis root meristem. Genes & Dev 17, 354–358.

Sarkar, A.K., Luijten, M., Miyashima, S., Lenhard, M., Hashimoto, T., Nakajima, K., Scheres, B., Heidstra, R., and Laux, T. (2007). Conserved factors regulate signalling in Arabidopsis thaliana shoot and root stem cell organizers. Nature 446, 811–814.

Shay, J.W., and Wright, W.E. (2019). Telomeres and telomerase: three decades of progress. Nat Rev Genet 20, 299–309.

Shimotohno, A., Heidstra, R., Blilou, I., and Scheres, B. (2018). Root stem cell niche organizer specification by molecular convergence of PLETHORA and SCARECROW transcription factor modules. Genes & Dev 32, 1085–1100.

Song, X., Leehy, K., Warrington, R.T., Lamb, J.C., Surovtseva, Y.V., and Shippen, D.E. (2008). STN1 protects chromosome ends in Arabidopsis thaliana. Proc Natl Acad Sci USA 105, 19815–19820.

Sozzani, R., Cui, H., Moreno-Risueno, M.A., Busch, W., Van Norman, J.M., Vernoux, T., Brady, S.M., Dewitte, W., Murray, J.A., and Benfey, P.N. (2010). Spatiotemporal regulation of cell-cycle genes by SHORTROOT links patterning and growth. Nature 466, 128–132.

Surovtseva, Y.V., Churikov, D., Boltz, K.A., Song, X., Lamb, J.C., Warrington, R., Leehy, K., Heacock, M., Price, C.M., and Shippen, D.E. (2009). Conserved telomere maintenance component 1 interacts with STN1 and maintains chromosome ends in higher eukaryotes. Mol Cell 36, 207–218.

van den Berg, C., Willemsen, V., Hendriks, G., Weisbeek, P., and Scheres, B. (1997). Short-range control of cell differentiation in the Arabidopsis root meristem. Nature 390, 287–289.

Walter, M., Chaban, C., Schutze, K., Batistic, O., Weckermann, K., Nake, C., Blazevic, D., Grefen, C., Schumacher, K., Oecking, C., et al. (2004). Visualization of protein interactions in living plant cells using bimolecular fluorescence complementation. Plant J 40, 428–438.

Weigel, D., and Glazebrook, J. (2002). Arabidopsis: A Laboratory Manual.

Welch, D., Hassan, H., Blilou, I., Immink, R., Heidstra, R., and Scheres, B. (2007). Arabidopsis JACKDAW and MAGPIE zinc finger proteins delimit asymmetric cell division and stabilize tissue boundaries by restricting SHORT-ROOT action. Genes & Dev 21, 2196–2204.

Yoo, S.D., Cho, Y.H., and Sheen, J. (2007). Arabidopsis mesophyll protoplasts: a versatile cell system for transient gene expression analysis. Nat Protoc 2, 1565–1572.

